# Pharmacogenetic and whole-brain activity analyses uncover integration of distinct molecular and circuit programs that drive learning

**DOI:** 10.1101/2022.02.05.479242

**Authors:** Jessica C. Nelson, Hannah M. Shoenhard, Michael Granato

**Affiliations:** Department of Cell and Developmental Biology, Perelman School of Medicine, University of Pennsylvania, Philadelphia, Pennsylvania, USA; Department of Cell and Developmental Biology; University of Colorado Anschutz Medical Campus, Aurora, CO, USA

**Keywords:** habituation learning, plasticity, zebrafish

## Abstract

Habituation is a foundational learning process critical for animals to adapt their behavior to changes in their sensory environment. Although habituation is considered a simple form of learning, the identification of a multitude of molecular pathways including several neurotransmitter systems that regulate this process suggests an unexpected level of complexity. How the vertebrate brain integrates these various pathways to accomplish habituation learning, whether they act independently or intersect with one another, and whether they act via divergent or overlapping neural circuits has remained unclear. To address these questions, we combined pharmacogenetic pathway analysis with unbiased whole-brain activity mapping using the larval zebrafish. This approach revealed five distinct molecular and circuit modules that regulate habituation learning and identified a set of molecularly defined brain regions associated with four of the five modules. Moreover, we find that in module 1 the palmitoyltransferase Hip14 cooperates with dopamine and NMDA signaling to drive plasticity, while in module 3 the adaptor protein complex subunit Ap2s1 drives habituation by antagonizing dopamine signaling, revealing two distinct and opposing roles for dopaminergic neuromodulation in the regulation of learning. Combined, our results define a core set of distinct modules that act in concert to regulate learning-associated plasticity, and provide compelling evidence that even seemingly simple learning behaviors in a compact vertebrate brain are regulated by a complex and overlapping set of molecular and circuit mechanisms.

## Introduction

Learning enables animals to modify their responses to stimuli based on prior experience. One of the simplest forms of learning is a non-associative plasticity mechanism termed habituation, which is defined by a gradual decrease in responding to repeated stimuli^1–3^. Habituation represents a foundation for more complex forms of plasticity and is observed in all animals. Habituation learning is also a pervasive feature of the nervous system, regulating response rates to stimuli spanning sensory modalities and including complex responses such as fear responses and feeding^4–6^. We previously established larval zebrafish as a model to study short term habituation learning^7^. In response to a sudden acoustic stimulus zebrafish perform a stereotyped acoustic startle response (ASR), comprised of a short-latency C-bend (SLC) escape response regulated by well-described hindbrain circuitry^8–10^. Repeated acoustic stimuli modulate sensory thresholds and result in habituation learning characterized by a gradual decline in response frequency^7,11–14^. Although this learning process appears simple at first glance, previous work revealed that at least long-term habituation learning is regulated by multiple mechanisms that operate on distinct time scales^2,15^. Similarly, numerous molecular-genetic mechanisms also regulate short-term habituation learning^16–26^. Moreover, pharmacological screens have identified multiple neurotransmitter and neuromodulatory systems contributing to habituation^7^. Despite their known relevance for learning, how individual habituation-regulatory pathways relate to one another, whether they act sequentially or regulate plasticity in parallel, and whether they are distributed over multiple brain areas or function within a common circuit is unclear.

Here, we first employed a pharmacogenetic approach to determine whether individual molecular regulators of habituation learning can be modulated by habituation-relevant neurotransmitter systems. To complement this pharmacogenetic approach, we then performed unbiased whole-brain imaging to define activity signatures for each pharmacogenetic manipulation, and to identify candidate brain regions in which habituation-regulatory modules exert their function. We identify five distinct molecular-circuit modules that regulate learning. Module 1 consists of the palmitoyl-transferase Hip14, as well as NMDA and dopamine signaling, while module 2 consists of Hip14 and one of its identified substrates, the voltage-gated Potassium channel subunit Kv1.1. Module 3 consists of the *pregnancy-associated plasma protein A* (PAPP-AA) and the AP2 adaptor complex subunit AP2S1, which together act to oppose dopamine signaling. Glycine signaling constitutes module 4, and the *voltage dependent calcium channel alpha2/delta subunit 3* gene (*cacna2d3*) defines module 5. Two of these modules reveal a critical role for dopamine signaling in the bi-directional modulation of habituation learning. Specifically, while the palmitoyltransferase Hip14 cooperates with neurotransmitter signaling through NMDA and dopamine receptors to drive habituation (module 1), the AP2 adaptor complex subunit AP2S1 promotes learning by opposing dopamine signaling (module 3). Moreover, while we find that three of the habituation-regulatory modules intersect mechanistically (modules 1, 2 and 3), modules 4 and 5 appear functionally independent from each other and from the other three interconnected modules, suggesting multiple habituation-regulatory mechanisms that act in parallel. Taken together, our findings highlight the strength of an integrative approach combining genetic and pharmacological manipulation of habituation learning with unbiased whole-brain activity mapping and reveal a more complete picture of the molecular and circuit mechanisms that drive vertebrate habituation learning.

## Results

### A sensitized habituation assay to uncover pharmacogenetic interactions

From an unbiased genetic screen we previously identified a set of five genes required for habituation learning^21,24,27,28^. To determine whether the identified molecular and circuit mechanisms regulate habituation learning independently of each other, or whether these genetic mechanisms converge at a common bottleneck, we set out to perform pathway analysis by exposing habituation mutants to pharmacological inhibitors of habituation-regulatory neurotransmitter signaling pathways. We reasoned that genetic and pharmacological manipulations that impinge upon components of independent or parallel pathways would enhance learning deficits while multiple insults to components of a common pathway would fail to produce additive deficits. However, impeding our ability to perform such analyses, we observed that genetic mutations that affect habituation learning, such as presumptive null mutations in the *zinc finger DHHC-type palmitoyltransferase gene zdhhc17*, encoding the palmitoyltransferase Hip14, result in a near complete loss of habituation at our standard stimulus intensities of 35.1dB (Figure 1A-B)^24^. This ceiling effect interferes with the ability to detect additive learning deficits, preventing us from detecting a further reduction in learning and thus preventing us from interpreting the results of the proposed pharmacogenetic pathway analysis. We therefore wondered whether reducing stimulus intensity might provide a sensitized assay in which mutant animals retain some capacity for habituation, and application of a pharmacological inhibitor of learning might reveal more severe habituation deficits. Consistent with previous findings that habituation learning is modulated by stimulus intensity^2^, we find that although still impaired relative to their siblings, *hip14* mutant animals are capable of habituation learning under conditions of reduced stimulus intensity (i.e. 0.4dB-25.6dB, Figure 1C-G). Moreover, these data reveal that presumptive null mutations in *hip14* fail to fully abolish habituation, and that when presented with lower intensity stimuli *hip14* mutant animals are capable of learning, albeit at a reduced level relative to their siblings. We conclude that at lower intensity, further learning impairments in *hip14* mutants induced by pharmacological inhibitors of learning might be readily detectable. We therefore selected 19.8dB for our sensitized learning assay and utilized this stimulus intensity to test five genetic mutants in combination with individual inhibitors of three neurotransmitter systems.

**Fig. 1.**
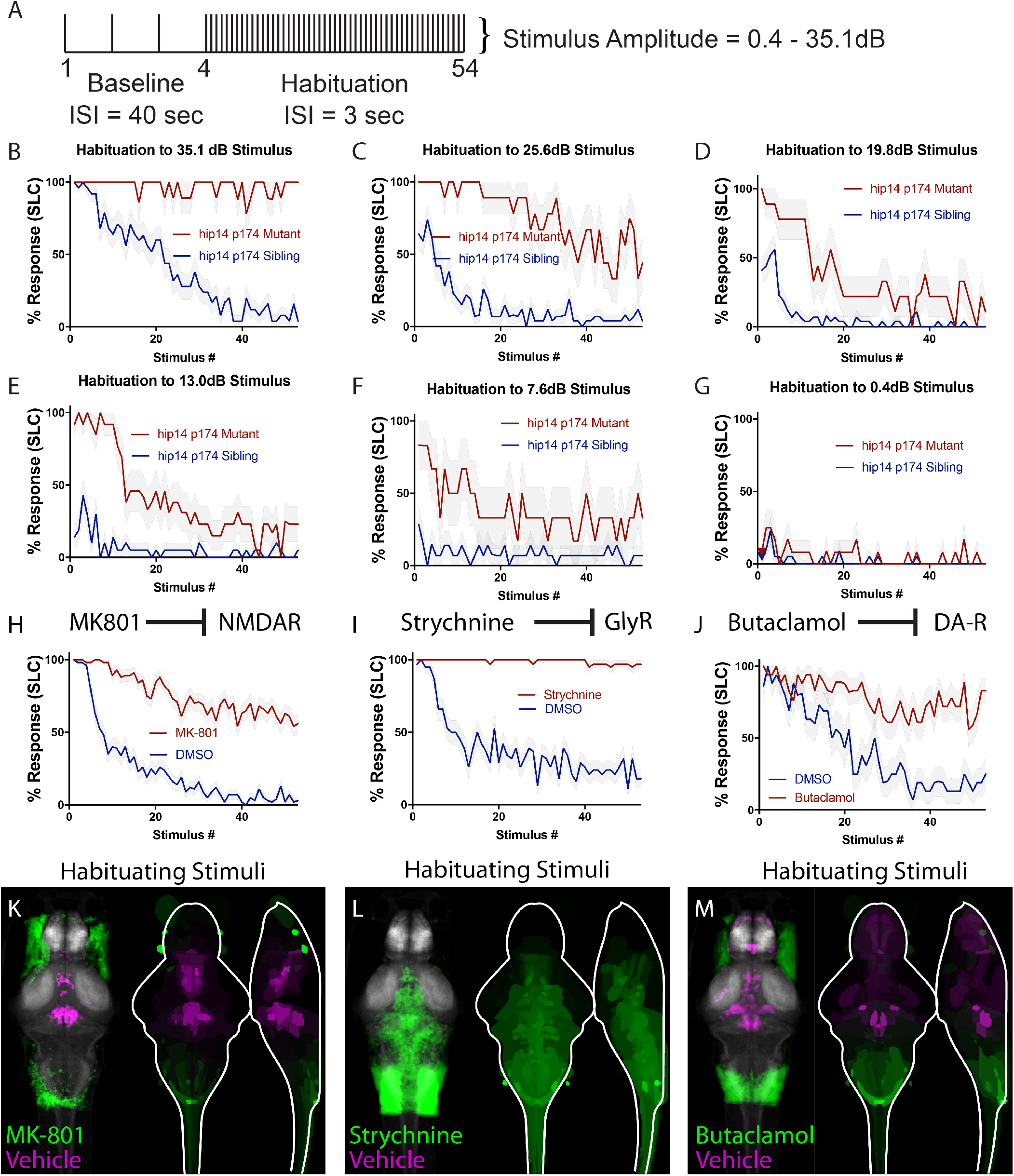
Sensitized learning assay and unbiased whole-brain imaging to examine impact of pharmacological inhibitors of habituation learning. **(A)** Stimulus paradigm used. ISI for baseline phase is 40 seconds, ISI for habituation phase is 3 seconds. **(B)** *hip14* mutants exhibit a complete failure to habituate to 35.1 dB acoustic stimuli. **(C-G)** Reduction in stimulus intensity as indicated results in a gradual increase in the ability of *hip14* mutants to habituate. **(H)** MK-801 is an NMDA inhibitor that strongly reduces habituation learning in 5-day old zebrafish larvae (n=58 DMSO-treated, n=57 MK-801-treated, stimulus intensity = 35.1dB). **(I)** Strychnine is a glycine receptor antagonist that strongly reduces habituation learning in 5-day old zebrafish larvae (n=38 DMSO-treated, n=38 Strychnine-treated, stimulus intensity = 35.1dB). **(J)** Butaclamol is a dopamine inhibitor that strongly reduces habituation learning in 5-day old zebrafish larvae (n=16 DMSO-treated, n=18 Butaclamol-treated, stimulus intensity = 35.1 dB). **(K-M)** Regions upregulated by the specified drug treatment under “Habituating Stimuli” conditions are indicated in green; regions downregulated are indicated in magenta. In all images, the left panel is a summed z-projection of the whole-brain activity changes. The right panel is a z-projection and x-projection of the analyzed MAP-map. Molecular targets of pharmacological agents are indicated with diagrams above each column. See S1 for brain activity maps under “No Stimulus” condition and “Non-Habituating Stimuli” condition. Also see **Supplemental Table S1** for ROIs identified in the experiments presented as well as in an independent replicate of each drug condition.

### Pharmacological inhibitors of habituation learning produce distinct patterns of neuronal activity

For the pharmacogenetic pathway analysis we selected the NMDA receptor inhibitor MK-801, the glycine receptor inhibitor Strychnine, and the dopamine receptor inhibitor Butaclamol. As previously reported, in 5-day old larval zebrafish, application of MK-801 results in significant impairments in habituation learning (Figure 1H)^7,29^. Whereas vehicle-exposed animals rapidly learn to ignore repeated acoustic stimuli, i.e. habituate, animals exposed acutely to the NMDA inhibitor continue to respond at a high rate (they fail to habituate). Similar effects are observed when animals are exposed to Strychnine^29^, (Figure 1I) or Butaclamol^7^ (Figure 1J). Despite their similar effects on habituation learning, we hypothesized that given their regulation of different neurotransmitter systems, these pharmacological agents might regulate learning through distinct effects on neuronal activity. In order to broadly assess brain activity signatures associated with each pharmacological treatment, we performed unbiased whole-brain activity mapping (MAP-mapping)^30^ for each of the three pharmacological inhibitors under three different acoustic stimulation conditions: “No Stimuli,” “Non-Habituating Stimuli,” and “Habituating Stimuli.” We find that each pharmacological agent produces a distinct activity pattern (Figure 1K-M, S1A–F). We find furthermore that the distinct patterns of activity induced by each inhibitor of learning are largely maintained across stimulation conditions (Figure 1K–M, S1A–F, Supplemental Table S1). In particular, MK-801 suppresses activity within the subpallium, habenula, and hypothalamus (Figure 1K, S1A,D), Strychnine produces widespread hyperactivation (Figure 1L, S1B,E), and Butaclamol treatment results in hindbrain hyperactivation and forebrain and diencephalic suppression relative to vehicle controls (Figure 1M, S1C,F). Our results that differences between inhibitor conditions, but not stimulation conditions, were readily detected reflects the design of our experiments, which were optimized to detect differences between drug conditions at the expense of sensitivity to differences in stimulation conditions. Together, these data suggest two possibilities: either that the neurotransmitter systems that regulate learning impinge upon different sets of circuit loci, which separately regulate habituation learning, or that their effects on learning are mediated through the limited regions that show overlapping activity changes.

### Hip14 acts through NMDA and dopamine signaling and produces broad hyperactivity of neuronal circuits

Having identified a sensitized habituation assay, and having established that pharmacological inhibitors of learning impinge upon activity within distinct brain regions, we set out to perform our pharmacogenetic analysis to examine all possible interactions between three neurotransmitter signaling inhibitors and five habituation mutants. We first tested whether Hip14 and NMDA receptors act together to regulate habituation learning by exposing mutant and sibling animals to either vehicle (DMSO) or the NMDA inhibitor MK-801 and then performing the sensitized habituation assay (Figure 1A,D). We found that while MK-801 severely reduced habituation in sibling animals, the same pharmacological manipulation in *hip14* mutant animals did not further reduce habituation learning (Figure 2A). This is consistent with a model in which Hip14 and NMDA act in a common pathway to drive habituation learning. When we plotted response frequency versus stimulus number in order to analyze the kinetics of learning (learning curves), we found that compared to MK-801-treated mutants, sibling animals treated with MK-801 exhibited more severe learning deficits (Figure 2B). This raises the possibility that compensatory, NMDA-independent learning mechanisms may be upregulated in *hip14* mutant animals. When we performed the same sensitized protocol in the presence of the glycine-receptor inhibitor Strychnine, we observed further reductions in learning in both siblings and *hip14* mutants, consistent with independent roles for glycine and Hip14 in the regulation of learning (Figure 2C-D). Finally, we performed our sensitized learning assay in the context of the dopamine receptor inhibitor Butaclamol. Here we found that dopamine receptor inhibition significantly impaired learning in sibling animals. However, the same manipulation did not reduce habituation learning in *hip14* mutant animals (Figure 2E-F). Taken together, our data provide strong evidence that Hip14 acts in a common pathway with dopamine and NMDA receptor signaling yet independently of glycine receptor signaling to drive habituation learning.

**Fig. 2.**
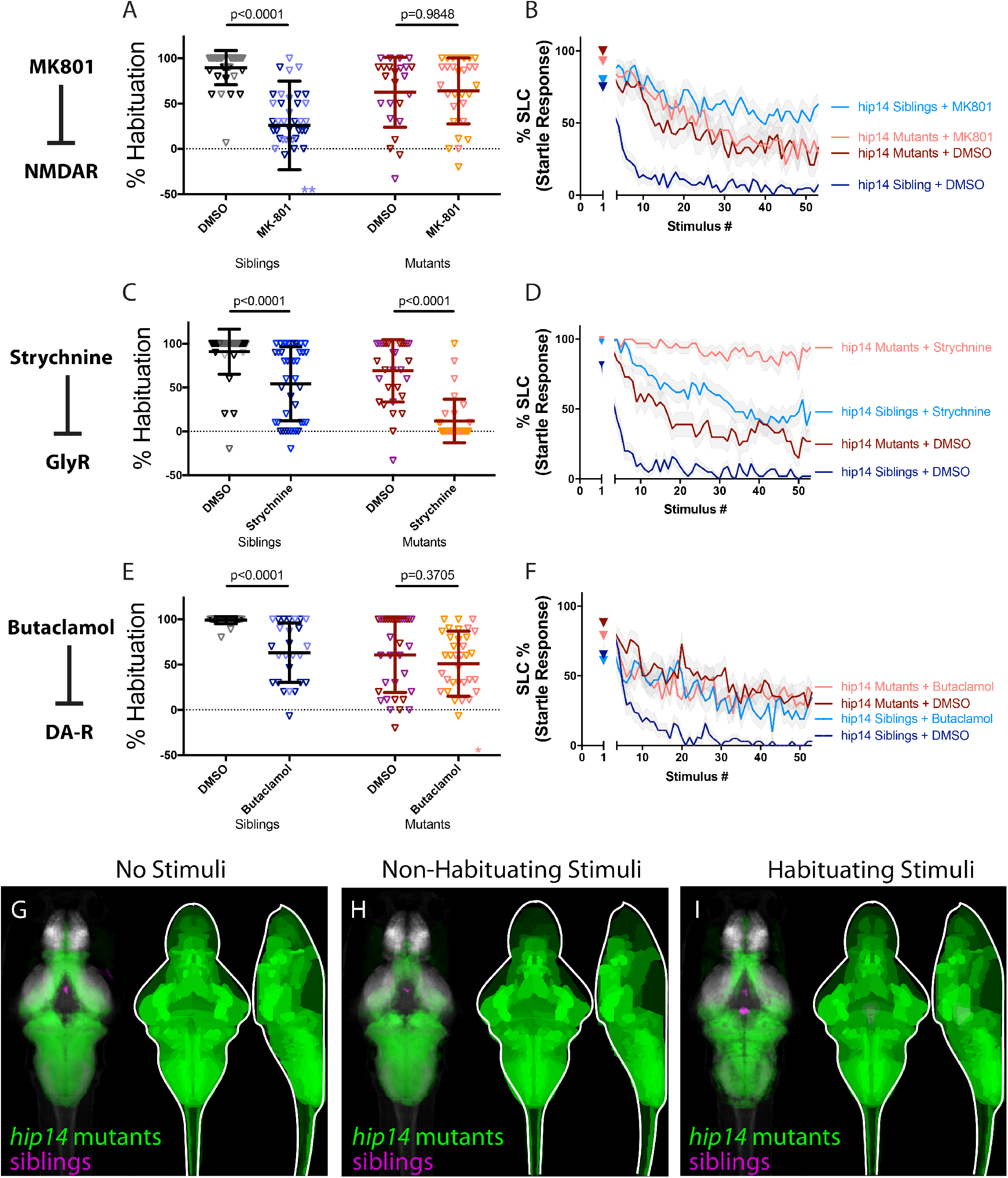
Hip14 acts through NMDA and dopamine signaling and produces broad hyperactivity of neuronal circuits. (**A-B**) MK-801 impairs learning in siblings, (p<0.0001 Sidak’s test for multiple comparisons, n=38 DMSO, n=38 MK-801), but does not enhance habituation learning deficits observed in *hip14* mutant larvae (p=0.9848, n=24 DMSO, n=29 MK-801, these and all subsequent statistical analyses use Sidak’s multiple comparisons test unless otherwise indicated). **indicate sibling+MK801 individuals with % hab values below y-axis limit at −100% and −180%. (**C-D**) Strychnine significantly enhances habituation learning deficits observed in *hip14* mutant larvae, indicating that glycine signaling and *hip14* may act within parallel molecular or circuit pathways to regulate learning (p<0.0001, n=40 DMSO-siblings, n=42 Strychnine-siblings, n=32 DMSO-mutants, n=32 Strychnine-mutants). (**E-F**) Butaclamol impairs learning in siblings (p<0.0001, n=33 DMSO, n=26 Butaclamol), but does not enhance habituation learning deficits observed in *hip14* mutant larvae (p=0.3705, n=34 DMSO, n=35 Butaclamol), indicating that dopamine receptor signaling and *hip14* may act within the same molecular or circuit pathway to regulate learning. *indicates a mutant+butaclamol individual with % hab value below y-axis limit at −60%. (**G-I**) Regions upregulated in *hip14* mutants are indicated in green; regions downregulated in *hip14* mutants are indicated in magenta. In all images, the left panel is a summed z-projection of the whole-brain activity changes. The right panel is a z-projection and x-projection of the analyzed MAP-map. Patterns of neuronal activity are similar between “No Stimuli” vs “Non-Habituating Stimuli” vs. “Habituating Stimuli” (restricted diencephalic downregulation of activity; nearly global upregulation of activity across the telencephalon, diencephalon, and rhombencephalon). See also **Supplemental Table S1** for ROIs up- and down-regulated in each condition, as well as in independent replicates.

In light of the finding that Hip14, dopamine, and NMDA signaling act in a common pathway to regulate habituation, we wondered whether *hip14* mutant animals and larvae treated with NMDA and dopamine inhibitors might display overlap in their activity signatures. To address this question, we performed whole-brain activity mapping in animals lacking *hip14*, analyzed the resultant MAP-maps, and compared them with those obtained from NMDA and dopamine inhibitor treated animals. In *hip14* mutant brains we observed broad hyperexcitability across the forebrain and hindbrain (Figure 2G-I). Despite this being a distinct pattern from that observed in NMDA- and dopamine-inhibited animals, we observed commonalities in the activity signatures. In particular, activity in the diencephalon was reduced, and a handful of rhombencephalic areas were upregulated by all three manipulations (Figure 2G-I, Supplemental Table S1). These overlapping activity changes, observed across multiple treatments, represent potential habituation-regulating loci through which these putative regulatory modules exert their function.

### Kv1.1 exhibits a unique activity signature and acts independently of NMDA, glycine, and dopamine signaling

We have previously shown that Hip14 acts in part through the voltage-gated Potassium channel subunit Kv1.1, which is encoded by the *potassium voltage-gated channel, shaker-related subfamily member 1a gene*, *kcna1a* ^24^. We performed pharmacogenetic pathway analysis in *kcna1a* mutants and found no evidence that Kv1.1 functions in a pathway with NMDA receptor signaling. Rather, the NMDA receptor inhibitor MK-801 induced significant learning deficits in both *kcna1a* siblings and mutants (Figure 3A-B). Similarly, glycine receptor inhibition induced significant learning deficits in both *kcna1a* mutants and siblings (Figure 3C-D). Finally, we observed significant enhancement of learning deficits through inhibition of dopamine signaling in both *kcna1a* siblings and mutants (Figure 3E-F). Taken together, these data are consistent with a model in which *kcna1a* acts independently of NMDA, dopamine, and glycine receptor signaling to regulate habituation learning.

**Fig. 3.**
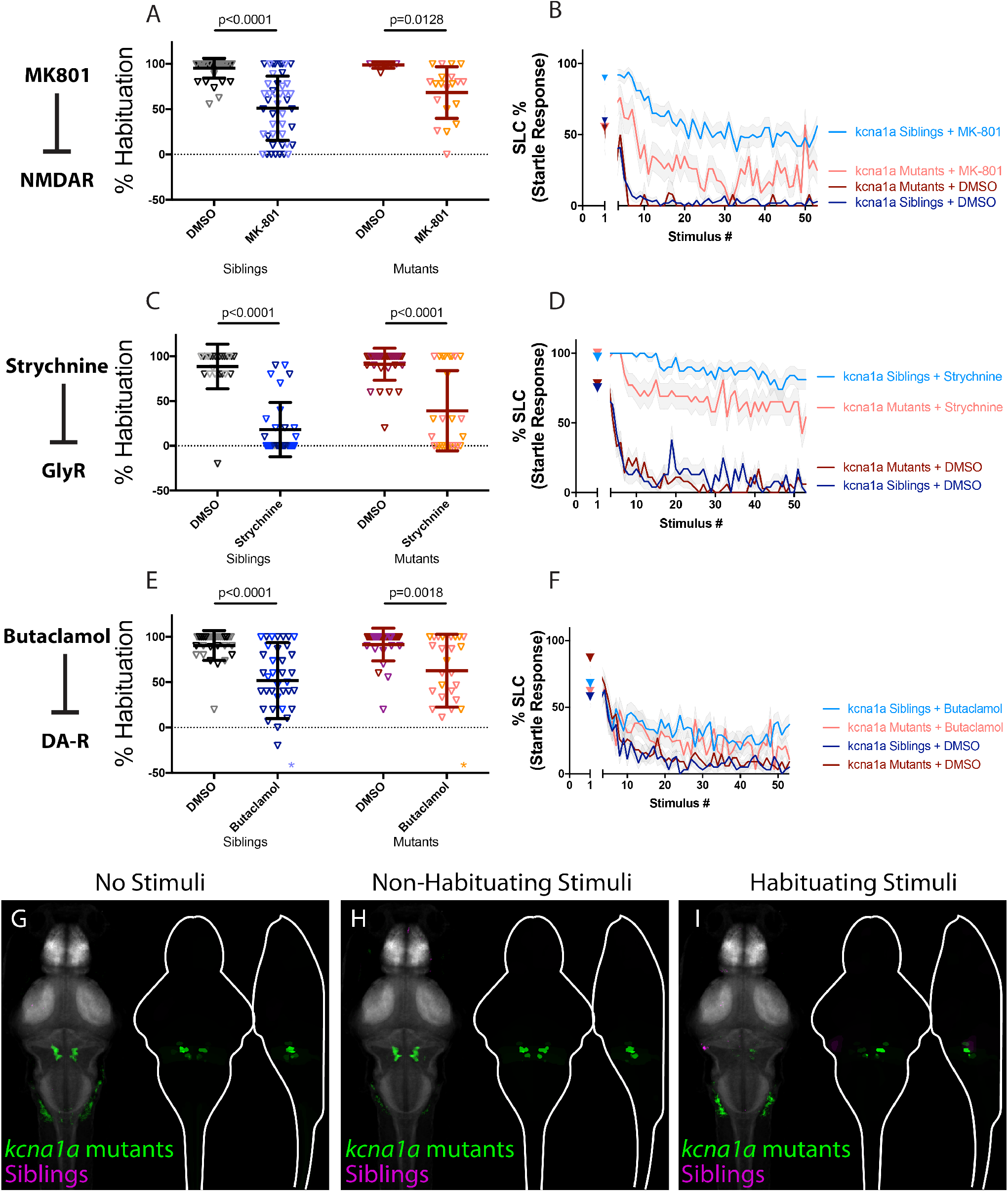
Kv1.1 exhibits a unique activity signature and acts independently of NMDA, glycine, and dopamine signaling. (**A-B**) MK-801 significantly enhances habituation learning deficits in *kcna1a* mutants and siblings (p<0.0001, n=42 DMSO-siblings, n=46 MK-801-siblings, p=0.0128, n=8 DMSO-mutants, n=20 MK-801 mutants). (**C-D**) Strychnine significantly enhances habituation learning deficits in *kcna1a* mutants and siblings (p<0.0001, n=24 DMSO-siblings, n=31 Strychnine-siblings, p<0.0001, n=35 DMSO-mutants, n=26 Strychnine-mutants). (**E-F**) Butaclamol significantly enhances habituation learning deficits in *kcna1a* mutants and siblings (p<0.0001, n=30 DMSO-siblings, n=37 Butaclamol-siblings, p=0.0018, n=30 DMSO-mutants, n=27 Butaclamol-mutants). *indicate a sibling+butaclamol and a mutant+butaclamol individual with % hab values below y-axis limit at −100% and −60% respectively. (**G-I**) Regions upregulated in *kcna1a* mutants are indicated in green; regions downregulated in *kcna1a* mutants are indicated in magenta. In all images, the left panel is a summed z-projection of the whole-brain activity changes. The right panel is a z-projection and x-projection of the analyzed MAP-map. Note the similar patterns of neuronal activity induced by “no stimuli” vs “non-habituating stimuli” vs. “habituating stimuli”: highly restricted upregulation of activity in the spiral fiber neuron clusters as well as in V2A (including Rom3) neurons. See also **Supplemental Table S1** for ROIs up- and down-regulated in each condition, as well as in independent replicates.

Next, we performed whole-brain activity mapping to identify where *kcna1a* might exert its function, and to assess whether its activity signature might overlap with that of other regulators of learning. We found a remarkably specific and unique pattern of activity induced by the loss of *kcna1a*. In particular, two populations known to express Kv1.1^24,31^ and involved in the execution of the escape response were found to be hyperactive: spiral fiber neurons and RoM3 excitatory reticulospinal (V2a) neurons (Figure 3G-I). We previously showed that Kv1.1 requires Hip14 for proper synaptic localization and hence likely acts downstream of Hip14 to regulate habituation learning. Consistent with these findings, we now find that the same populations that are hyperactive in *kcna1a* mutants are also hyperactive in *hip14* mutants in all conditions except for one (Non-Habituating Stimuli Replicate 2 of 3). Moreover, the observation that activity changes are more restricted in *kcna1a* mutant brains compared to those observed in *hip14* mutant brains is consistent with our prior observation of more severe learning deficits in *hip14* mutants as compared to *kcna1a*^24^. These data lend further support to our hypothesis that Hip14 acts through other substrates besides Kv1.1 to regulate habituation. Combining these results with the findings of our pharmacogenetic analysis, we conclude that Kv1.1 likely functions in a restricted set of hindbrain neurons to carry out NMDA- and dopamine-independent functions downstream from Hip14.

### PAPP-AA promotes habituation by limiting endogenous dopamine signaling

The previous genetic screen^21^ additionally identified the *pregnancy-associated plasma protein A* (*pappaa*) gene as a critical regulator of habituation learning. PAPP-AA has been shown to act through regulation of Insulin Growth Factor Receptor (IGFR) signaling to regulate learning^21^, yet it is not known whether PAPP-AA interacts with any of the other identified habituation regulatory pathways. In order to investigate this question, we performed our pharmacogenetic pathway analysis in *pappaa* mutants. Compared to DMSO treated *pappaa* mutants, application of MK-801 or Strychnine to mutant animals resulted in further reduction of habituation learning, providing compelling evidence that PAPP-AA promotes habituation learning independent of NMDA (Figure 4A-B), and glycine receptor signaling (Figure 4C-D). In contrast, treatment of *pappaa* mutants with the dopamine receptor antagonist Butaclamol failed to enhance learning deficits in *pappaa* mutants when compared to DMSO treated mutants, and in fact trended toward ameliorating learning deficits in *pappaa* mutants (p=0.0731) (Figure 4E-F)). These data suggest that PAPP-AA may be required to suppress dopamine signaling and are consistent with a scenario in which dopaminergic inhibition somewhat normalizes behavioral deficits in *pappaa* mutants.

**Fig. 4.**
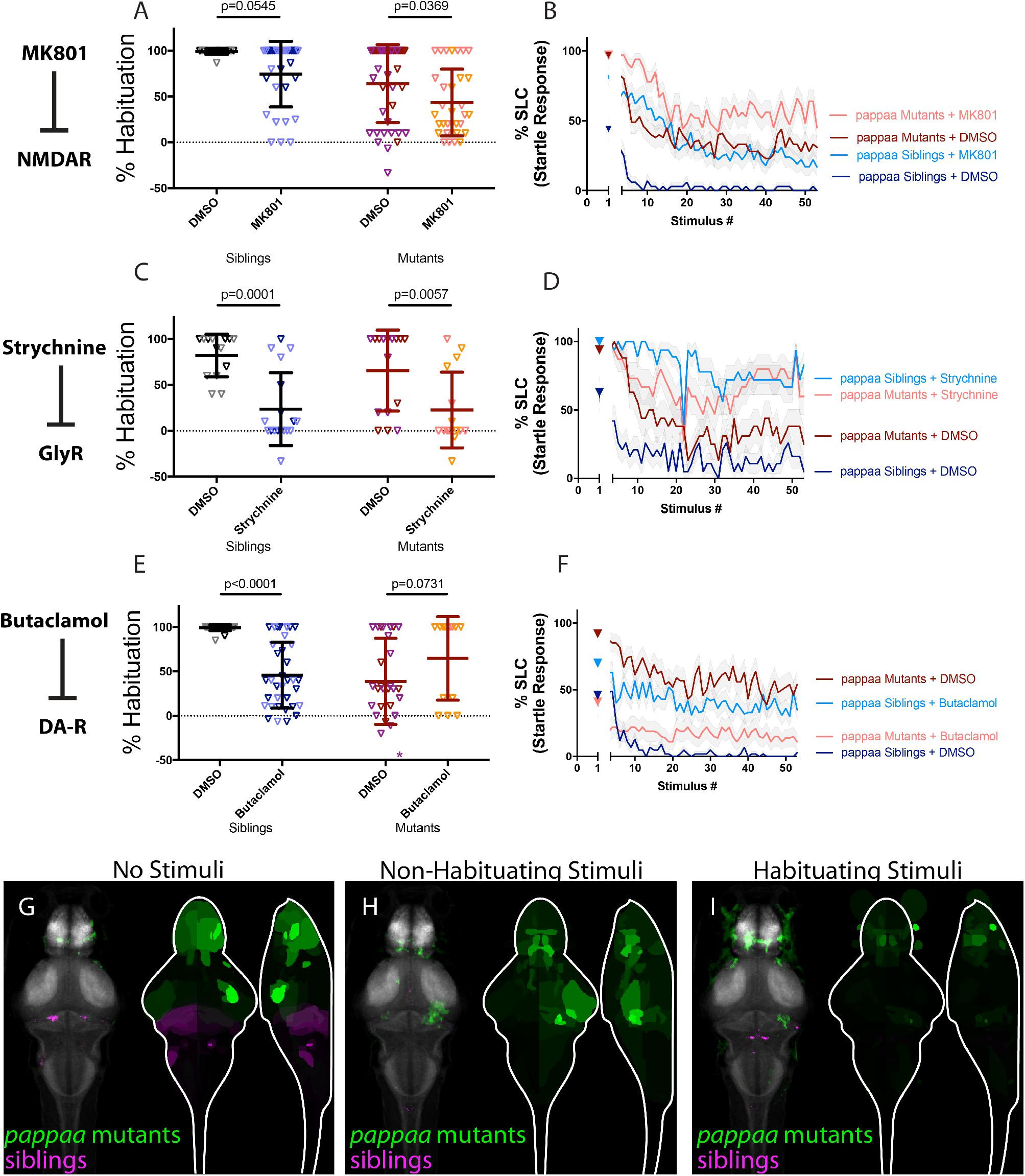
PAPP-AA promotes habituation by limiting endogenous dopamine signaling. (**A-B**) MK-801 significantly enhances habituation learning deficits observed in *pappaa* mutant larvae, indicating that NMDA signaling and *pappaa* may act within parallel molecular or circuit pathways to regulate learning (p=0.0369, n=39 DMSO-mutants, n=32 MK-801 mutants; p=0.0545, n=16 DMSO-siblings, n=31 MK-801 siblings). (**C-D**) Strychnine significantly enhances habituation learning deficits observed in *pappaa* mutant larvae, indicating that glycine signaling and *pappaa* may act within parallel molecular or circuit pathways to regulate learning (p=0.0057, n=16 DMSO-mutants, n=15 Strychnine-mutants; p=0.0001, n=14 DMSO-Siblings, n=18 Strychnine-siblings). (**E-F**) Butaclamol does not significantly enhance habituation learning deficits observed in *pappaa* mutant larvae, but rather trends toward significantly restoring learning (p<0.0001, n=31 DMSO-siblings, n=35 Butaclamol-siblings; p=0.0731, n=28 DMSO mutants, n=13 Butaclamol mutants). *indicates a mutant+DMSO individual with % hab value below y-axis limit at −100% (**G-I**) Regions upregulated in *pappaa* mutants are indicated in green; regions downregulated in *pappaa* mutants are indicated in magenta. In all images, the left panel is a summed z-projection of the whole-brain activity changes. The right panel is a z-projection and x-projection of the analyzed MAP-map. Patterns of neuronal activity are similar between “no stimuli” vs “non-habituating stimuli” vs. “habituating stimuli” (increased activity within the telencephalon and hypothalamus; decreased activity within multiple rhombencephalic loci). Note that this pattern is somewhat inverted relative to that observed in Butaclamol-treated animals, consistent with a role for *pappaa* in regulating dopamine signaling. See also **Supplemental Table S1** for ROIs up- and down-regulated in each condition, as well as in independent replicates.

Finally, we performed whole-brain activity mapping in *pappaa* mutant animals. Upon analyzing the resultant MAP-maps, we observed a subtle downregulation of neuronal activity particularly in the hindbrain, as well as upregulation of activity particularly in the pallium, subpallium, and hypothalamus (Figure 4G-I). These activity patterns are inverted when compared to those obtained by treatment of wild type animals with the dopamine antagonist Butaclamol, where activity in the hindbrain is increased and activity within the pallium, subpallium, and hypothalamus are decreased. These opposing activity signatures in *pappaa* mutants and dopamine-inhibited animals, together with the finding that Butaclamol restores learning in *pappaa* mutants, are consistent with a scenario in which PAPP-AA regulates habituation learning by limiting endogenous dopamine signaling.

### CACNA2D3 acts independently of other regulators of habituation learning

We recently identified the *calcium channel voltage dependent alpha2/delta subunit 3* gene, *cacna2d3*, encoding an auxiliary subunit of the voltage-gated calcium channel (VGCC) complex, as a genetic regulator of habituation learning^28^. We wondered how this VGCC subunit cooperates with the other regulators of habituation learning and therefore repeated our pharmacogenetic screen in the *cacna2d3* mutant background. We found that like *pappaa* and *kcna1a*, treatment of *cacna2d3* mutants with either the NMDA inhibitor MK801 or the glycinergic signaling inhibitor Strychnine further reduced habituation learning when compared to DMSO treated *cacna2d3* mutants (Figure 5A-D), consistent with a model in which *cacna2d3* regulates habituation independently of NMDA and glycinergic signaling. We next examined the interaction between *cacna2d3* and dopamine signaling (Figure 5E). Analysis of the learning curves for this experiment revealed an almost flat learning curve for Butaclamol-treated siblings (Figure 5F), consistent with a strong effect of dopaminergic inhibition on learning in *cacna2d3* mutants, and consistent with a model in which dopamine and *cacna2d3* function in parallel to regulate learning. Although this effect did not reach statistical significance, the impact of Butaclamol on *cacna2d3* mutant learning curves suggests that *cacna2d3* functions independently from NMDA, glycine, and dopamine signaling.

**Fig. 5.**
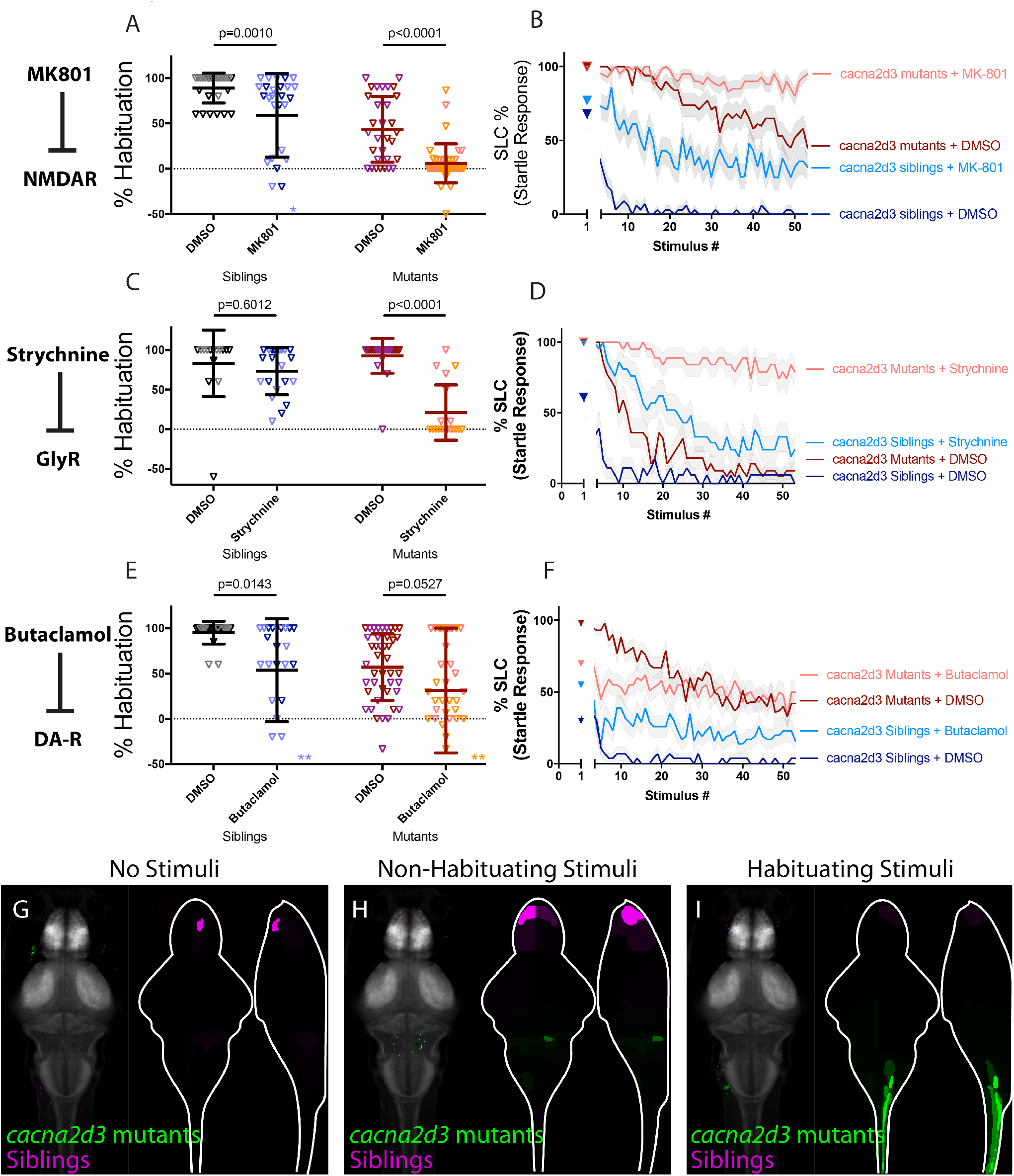
CACNA2D3 acts independently of other regulators of habituation learning. (**A-B**) MK-801 significantly enhances habituation learning deficits observed in *cacna2d3* mutant larvae, indicating that NMDA signaling and *cacna2d3* may act within parallel molecular or circuit pathways to regulate learning (p=0.0010, n=28 DMSO siblings, n=27 MK-801 siblings, vs. p<0.0001, n=31 DMSO mutants, n=41 MK-801 mutants). *indicates a sibling+MK801 individual with a % hab value below y-axis limit at −60% (**C-D**) Strychnine significantly enhances habituation learning deficits observed in *cacna2d3* mutant larvae, indicating that glycine signaling and *cacna2d3* may act within parallel molecular or circuit pathways to regulate learning (p=0.6012, n=15 DMSO siblings, n=21 Strychnine siblings vs. p<0.0001, n=22 DMSO mutants, n=19 Strychnine mutants). (**E-F**) Butaclamol does not significantly enhance habituation learning deficits observed in *cacna2d3* mutant larvae (p=0.0143, n=20 DMSO siblings, n=24 Butaclamol siblings, vs. p=0.0527, n=43 DMSO mutants, n=35 Butaclamol mutants). However, inspection of the learning curves (**F**) reveals a dramatic difference in the learning curves of mutants with or without drug. *indicate 2 sibling+Butaclamol and 2 mutant + butaclamol individuals with % hab value below y-axis limit at −100%, −60%, −256%, and −80% respectively. (**G-I**) Regions upregulated in *cacna2d3* mutants are indicated in green; regions downregulated in *cacna2d3* mutants are indicated in magenta. In all images, the left panel is a summed z-projection of the whole-brain activity changes. The right panel is a z-projection and x-projection of the analyzed MAP-map. Unlike other mutants, *cacna2d3* mutants do not exhibit reproducible changes in neuronal activity relative to their siblings in any stimulation condition. See also **Supplemental Table S1** for ROIs up- and down-regulated in each condition, as well as in independent replicates.

When we performed whole-brain imaging in *cacna2d3* mutants, we observed inconsistent activity changes across all stimulus conditions (Figures 5G-I). This lack of a defined whole-brain activity signature is unique to *cacna2d3* among the three pharmacological and five genetic manipulations that we tested. We interpret these results to reflect that CACNA2D3 may induce only subtle changes in neuronal activity or that it may simultaneously upregulate and downregulate activity within physically commingled neuronal populations.

### AP2S1 promotes habituation by limiting endogenous dopamine signaling

We recently identified a splice site mutation in the *adaptor related protein complex 2 subunit sigma 1* (*ap2s1*) gene, positioning the AP-2 adaptor complex as a fifth genetic regulator of habituation learning. We previously demonstrated that besides its role in habituation learning, AP-2 modulates sensorimotor decision-making via the Calcium-Sensing Receptor, CaSR^27^. Yet whether AP-2 also modulates NMDA, dopamine, or glycinergic signaling to regulate habituation learning has not been examined. We therefore performed our sensitized learning assay in *ap2s1* mutants and siblings and found that NMDA-receptor inhibition by MK-801 significantly impaired learning in *ap2s1* mutants, indicating that these two regulators of learning function in parallel (Figure 6A-B). Although not statistically significant (p=0.0773), glycinergic inhibition in the context of *ap2s1* mutations also revealed a clear and dramatic trend toward enhancement of learning deficits (Figure 6C-D). Finally, while Butaclamol-mediated inhibition of dopamine signaling led to significantly impaired learning in *ap2s1* siblings, Butaclamol treatment of *ap2s1* mutants significantly restored learning (p=0.0206; Figure 6E-F).

**Fig. 6.**
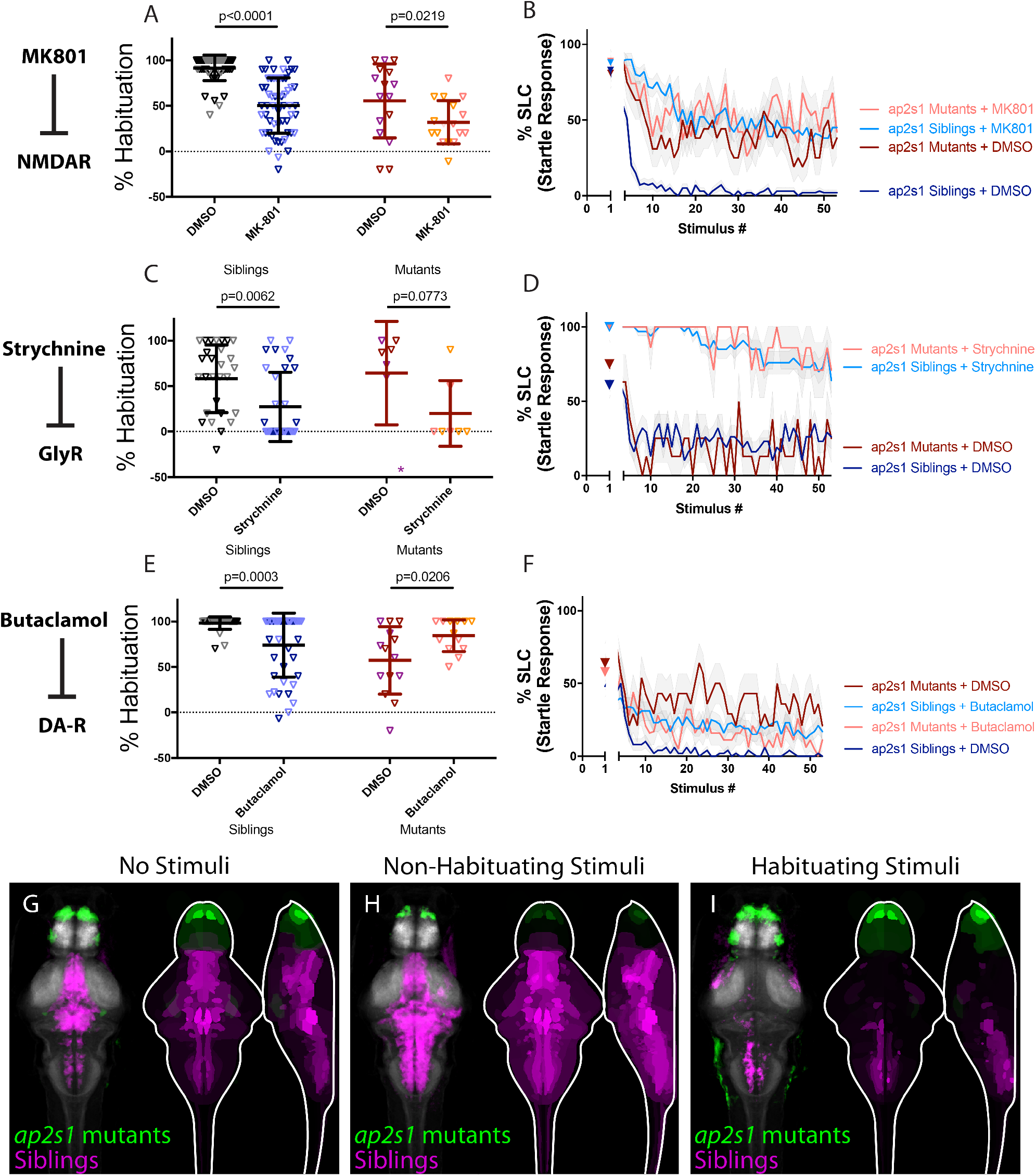
AP2S1 promotes habituation by limiting endogenous dopamine signaling. (**A-B**) MK-801 significantly enhances habituation learning deficits observed in *ap2s1* mutant larvae, indicating that NMDA signaling and *ap2s1* may act within parallel molecular or circuit pathways to regulate learning (p<0.0001, n=54 DMSO Siblings, n=58 MK-801 siblings; p=0.0219, n=16 DMSO mutants, n=17 MK-801 mutants). (**C-D**) Strychnine trends toward enhancing habituation learning deficits observed in *ap2s1* mutant larvae (p=0.0062, n=29 DMSO siblings, n=33 Strychnine siblings; p=0.0773, n=7 DMSO mutants, n=7 Strychnine mutants). Inspection of learning curves in (**D**) shows dramatic differences between sibling and *ap2s1* mutant larvae, indicating that glycine signaling and *ap2s1* act within parallel molecular or circuit pathways to regulate learning. *indicates a mutant+DMSO individual with a % hab value below y-axis limit at −60% (**E-F**) While Butaclamol inhibits learning in siblings (p=0.0003, n=36 DMSO siblings, n=42 Butaclamol siblings) it does not significantly enhance habituation learning deficits in *ap2s1* mutant larvae. Rather, *ap2s1* mutant animals learn significantly more robustly in the presence of the normally habituation-blocking Butaclamol (p=0.0206, n=14 DMSO mutants, n=13 Butaclamol mutants). (**G-I**) Regions upregulated in *ap2s1* mutants are indicated in green; regions downregulated in *ap2s1* mutants are indicated in magenta. In all images, the left panel is a summed z-projection of the whole-brain activity changes. The right panel is a z-projection and x-projection of the analyzed MAP-map. Consistent with the even stronger effect of Butaclamol in driving learning in *ap2s1* mutants relative to *pappaa* mutants, *ap2s1* mutant animals exhibit an even more dramatically inverted pattern relative to Butaclamol-treated animals. *ap2s1* mutant animals exhibit robust upregulation in the telencephalon (while Butaclamol-treated animals show downregulation here). Similarly, *ap2s1* mutants show dramatically downregulated activity within the rhombencephalon, while our Butaclamol results indicate that dopamine inhibition upregulates activity here. See also **Supplemental Table S1** for ROIs up- and down-regulated in each condition, as well as in independent replicates.

Finally, we performed whole-brain activity mapping in *ap2s1* mutants. Given that inhibition of dopamine partially restored habituation in both *pappaa* and *ap2s1* mutants, we predicted that *ap2s1* mutants would exhibit a similar activity pattern to that observed in *pappaa* mutants, and an inverted pattern with respect to dopamine receptor-inhibited animals. Indeed, analysis of whole-brain activity maps in *ap2s1* mutants revealed activity patterns similar to those we observed in *pappaa* mutants (Figure 6G-I), characterized by a marked downregulation in areas of the hindbrain that were observed to be upregulated in Butaclamol-treated animals, including a small hindbrain cluster of Tyrosine Hydroxylase (th, the enzyme required for dopamine synthesis) positive neurons (**Supplemental Table S1**). Moreover, we noted significant upregulation of activity in the subpallium, pallium, and intermediate hypothalamus, all areas that saw significant upregulation in *pappaa* mutant brains and downregulation in Butaclamol-treated animals (**Supplemental Table S1**). Combined, these results suggest a model in which AP2S1, like PAPP-AA, is involved in the suppression of dopamine signaling. As in the case of *pappaa*, loss of *ap2s1* results in a dysregulation of dopaminergic signaling that can be restored through its pharmacological inhibition via Butaclamol.

In summary, comparing pharmacogenetic analyses and brain activity signatures across five different habituation genes and three inhibitors of habituation-regulatory neurotransmitter pathways reveals distinct molecular and circuit modules that regulate habituation learning and identifies molecularly defined brain regions associated with each of the modules.

## Discussion

We set out to map genetic regulators onto the circuit / neuro-transmitter systems that drive habituation learning. We employed two complementary strategies. First, we developed a sensitized habituation learning assay, utilizing an acoustic stimulus of intermediate intensity, in which additive learning deficits caused by combining genetic and pharmacological regulators of habituation can readily be detected and quantified. We reasoned that the habituation deficits caused by a given genetic mutation would be enhanced by pharmacological manipulations of independent or parallel habituation-regulatory pathways, but not by manipulation of habituation-regulatory mechanisms in the same pathway module. This approach has been utilized in previous studies in which manipulations that increase dopaminergic signaling render animals hypo-responsive to dopamine receptor agonism^32^. Similarly, manipulations that decrease NMDA receptor localization render animals hypo-responsive to NMDA receptor antagonism^33^.

Second, we performed MAP-mapping in the context of each pharmacological or genetic manipulation, resulting in a unique set of brain activity maps, all of which reflect habituation learning deficient patterns of activity. We are struck by the diversity of brain activity patterns associated with deficits in habituation learning. Although overlapping patterns were observed for *hip14*, MK-801, and Butaclamol, as well as for *pappaa* and *ap2s1*, we note that these two putative modules differ from one another, and from the patterns observed for Strychnine and *kcna1a*, as well as from the observed lack of activity changes in *cacna2d3* mutants. Taken together, these results are consistent with at least two potential interpretations. First, it is possible that although each perturbation broadly impacts brain activity in distinct ways, brain activity maps for regulators of habituation learning overlap within a handful of critical regions that drive learning. A second interpretation is that habituation learning is heavily regulated and involves the cooperation of multiple parallel genetic-circuit modules. The latter interpretation is consistent with our observation that some genetic regulators of habituation learning show a significant interaction with NMDA and/or dopamine signaling, while others do not. Moreover, the existence of parallel short-term habituation-regulatory modules mirrors the previous finding that long-term habituation learning in the larval zebrafish is regulated by multiple parallel processes^15^.

Our work reveals five habituation regulatory modules (Figure 7A). The first module consists of Hip14, as well as NMDA and dopamine signaling (Figure 7B). Our pharmacological screen uncovered significant pharmacogenetic interactions between *hip14* and inhibitors of both NMDA and dopamine receptor signaling. Additionally, an unbiased clustering algorithm identified that brain activity patterns produced by the NMDA inhibitor MK-801 and the dopamine receptor antagonist Butaclamol are similar, and the relative strengths of activity changes from these treatments are highly correlated (R-squared=0.71, p<0.05; S2A-B). Although both Strychnine and mutations in *hip14* broadly upregulate neuronal activity, the specific regions that they upregulate are only weakly correlated (R-squared=0.24, p<0.05, S2C). This finding is consistent with the results from our pharmacogenetic approach, which placed these two regulators into pathways independent of each other.

**Fig. 7.**
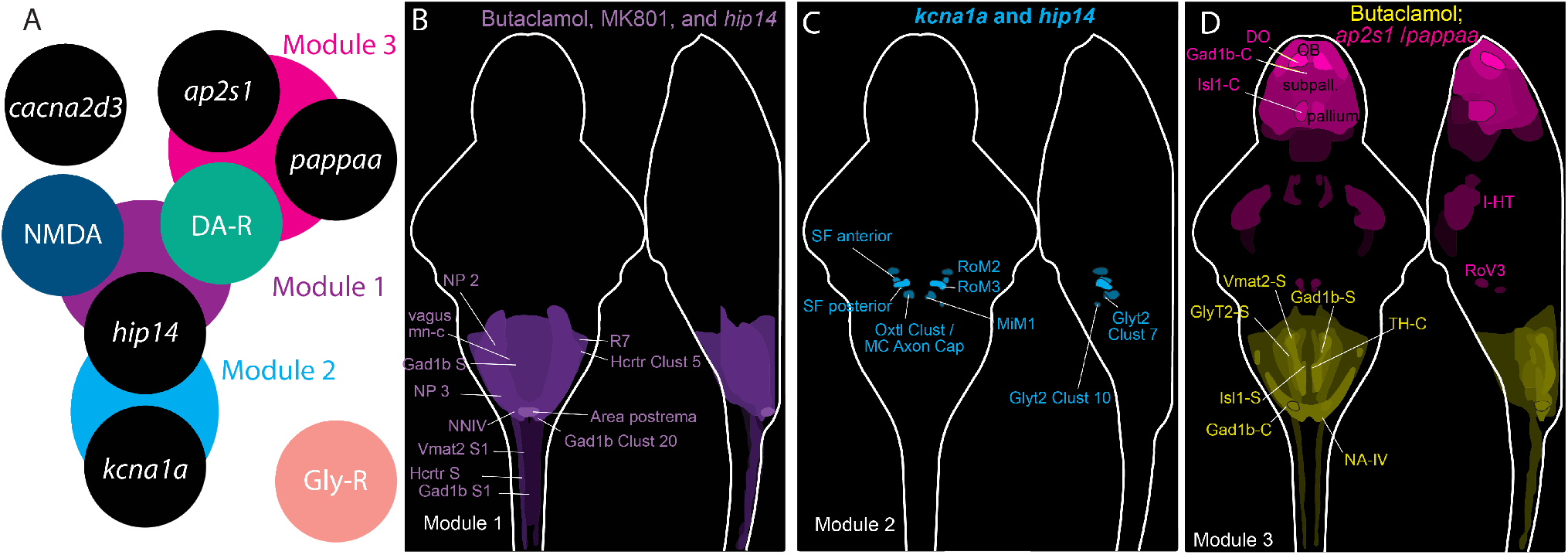
Cluster analysis identifies habituation regulatory modules. (**A**) Module 1 (purple) comprises *hip14*, NMDA, and dopamine receptors. Module 2 (aqua) comprises *hip14* and *kcna1a*. Module 3 (pink) comprises *ap2s1*, *pappaa*, and dopamine receptors. Modules 4 and 5 are comprised of *cacna2d3* and glycine receptor signaling acting in parallel to all other modules. (**B**) Regions commonly upregulated by dopamine inhibition, NMDA inhibition, and mutations in *hip14* are indicated in purple. (**C**) Regions upregulated by mutations in *kcna1a* and *hip14* are indicated in blue. (**D**) Regions downregulated by dopamine receptor inhibition and upregulated by mutations in *ap2s1* or *pappaa* are indicated in pink. Regions upregulated by dopamine receptor inhibition and downregulated by mutations in *pappaa* or *ap2s1* are indicated in yellow. See also S2 for cluster analysis heat map and correlations between pharmacogenetic treatments. In (**B-D**), signal intensity is proportional to the sum of the absolute intensity values. Abbreviations in (**B**): NP2 = Rhombencephalon – Neuropil Region 2, X vagus mn cluster = Rhombencephalon - X Vagus motorneuron cluster, Gad1b S = Rhombencephalon Gad1b Stripe 2, NP3 = Rhombencephalon Neuropil Region 3, NNIV = Rhombencephalon - Noradrendergic neurons of the Interfascicular and Vagal areas, Vmat2 S1 = Spinal Cord - Vmat2 Stripe1, Hcrtr S = Spinal Cord - 6.7FDhcrtR-Gal4 Stripe, Gad1b S1 = Spinal Cord - Gad1b Stripe 1, R7 = Rhombomere 7, Hcrtr Clust 5 = Rhombencephalon - 6.7FDhcrtR-Gal4 Cluster 5, Area postrema = Rhombencephalon Area Postrema, Gad1b Clust 20 = Rhombencephalon - Gad1b Cluster 20. Abbreviations in (**C**): SF anterior = Rhombencephalon - Spiral Fiber Neuron Anterior cluster, SF posterior = Rhombencephalon - Spiral Fiber Neuron Posterior cluster, Oxtl Clust MC Axon Cap = Rhombencephalon - Oxtl Cluster 2 Near MC axon cap, RoM2 = Rhombencephalon - RoM2, RoM3 = Rhombencephalon – RoM3, MiM1 = Rhombencephalon - MiM1. Abbreviations in (**D**): DO = Telencephalon - Olfactory bulb dopaminergic neuron areas, Gad1b-C = Telencephalon - Subpallial Gad1b cluster, Isl1-C = Telencephalon - Isl1 clusters 1 and 2, OB = Telencephalon - Olfactory Bulb, Subpall = Telencephalon – Subpallium, Pallium = Telencephalon – Pallium, Vmat2-S = Rhombencephalon - Vmat2 Stripe2, GlyT2-S = Rhombencephalon - Glyt2 Stripe 2, Isl1b-S = Rhombencephalon Isl1 Stripe1, Gad1b-C = Rhombencephalon - Gad1b Cluster 20, Gad1b-S = Rhombencephalon - Gad1b Stripe 2, TH-C = Rhombencephalon - Small cluster of TH stained neurons.

Our second module consists of Hip14 and Kv1.1 (Figure 7C). Mutations in both *hip14* and *kcna1a* strongly upregulate activity in the spiral fiber neuron clusters as well as in V2a neurons (S2D). Dysfunction of V2a neurons within the spinal cord was previously proposed as a potential mechanism underlying the kinematic deficits observed in *kcna1a* mutants^31^, and we now hypothesize that hyperactivation of V2a neurons within the hindbrain could contribute to the habituation deficits observed in *kcna1a* mutants. Spiral fibers also constitute an attractive locus for Kv1.1’s activity in regulating learning. We previously found that Hip14 can palmitoylate Kv1.1 and regulates its localization to the spiral fiber terminals^24^. Moreover spiral fibers are known to undergo plasticity during habituation learning^34^. Interestingly, these neurons were not reliably identified as showing differential activity in MK-801 or Butaclamol-treated animals (Module 1). It is possible that changes within spiral fibers are subtle enough to be missed by our MAP-mapping analysis. Alternatively, Hip14, NMDA, and dopamine signaling could regulate habituation learning in a pathway that is parallel to the role of Hip14 and Kv1.1 within spiral fibers. This explanation is consistent with *hip14*’s stronger habituation deficit relative to *kcna1a*, as well as the finding that Hip14 acts in a common pathway with dopaminergic and NMDA signaling while Kv1.1 does not.

The third module includes PAPP-AA and AP2S1 and is further defined by its unique relationship to dopamine signaling (Figure 7D). While Hip14 seems to promote dopaminergic signaling (impinging upon both pathways results in no enhancement of learning deficits), our data are consistent with PAPP-AA and AP2S1 opposing dopamine signaling (inhibition of signaling through dopamine receptors normalizes habituation learning in *ap2s1* and *pappaa* mutants). Moreover, our MAP-mapping analysis revealed regions whose activity is oppositely regulated by Butaclamol and *ap2s1*/*pappaa*, particularly in the telencephalon (upregulated by *ap2s1* and *pappaa* and downregulated by Butaclamol) and hindbrain (upregulated by Butaclamol and downregulated by *ap2s1*) (S2E-F). We hypothesize that the opposing function of D1 and D2/D3-type dopamine receptors in regulating the startle response may help to explain these surprising results^35^. While dopamine acts to drive habituation learning (reducing stimulus responsiveness) through D1-type dopamine receptors, it is known that it additionally promotes stimulus responsiveness through D2/D3-type dopamine receptors in mice^35^. We propose that AP2S1 and PAPP-AA are required to limit signaling through D2/D3-type dopamine receptors. In this scenario, mutations in either gene would result in excessive D2/D3 signaling and hyperresponsive larvae that fail to habituate to acoustic stimuli. Under these conditions, applying a dopamine receptor antagonist might normalize D2/D3 signaling and stimulus responsiveness, allowing animals to habituate. In support of these findings, work in zebrafish has found that high doses of the D2 receptor antagonist amisulpride can promote habituation learning^36^, and that the D2 antagonist haloperidol promotes long-term habituation of the O-bend or visual startle^15^.

We additionally note that the brain activity patterns observed for AP2S1 resemble the brain activity maps recently published for low-habituating populations obtained through breeding selection, showing increased neuronal activity in the forebrain, and decreased activity in the hindbrain^37^. Moreover, our data suggest that dopaminergic neuromodulation is an important driver of activity in the larval telencephalon and are supported by recent optogenetic experiments performed in Th2-expressing neurons^38^. However, in contrast to our findings, previous work found that ablating dopaminergic neurons of the caudal hypothalamus did not alter habituation and that dopamine neuron activity was actually elevated in high-habituating populations relative to low^37^. Our opposing results underscore the complex roles that dopamine plays in regulating acoustic startle sensitivity and habituation. Additionally, we note that chemogenetic ablation was restricted to Th1-positive dopamine neurons, and that this approach is chronic, in contrast to our transient pharmacological inhibition, and eliminates dopaminergic neurons while potentially having no immediate effect on neurotransmitter levels^39^. Given these differences, it is perhaps not surprising that these approaches yield somewhat differing results. Further work will be required to understand the role of dopamine in regulating habituation learning.

Finally, our work is consistent with glycine signaling (Module 4) and *cacna2d3* (Module 5) functioning in parallel to one another and to the other three modules described. We find no evidence that *cacna2d3* acts to regulate dopaminergic, glycinergic, or NMDA signaling, and likewise find no pharmacogenetic interactions between Strychnine and *hip14*, *kcna1a*, *pappaa*, or *ap2s1*. Although it is possible that a weak interaction between these and the other modules went undetected by our approach, we look forward to investigating potential interactions between these pathways and other pharmacogenetic regulators of learning.

One striking and unexpected finding that arose from our data is that each pharmacological manipulation or genetic mutation induced a brain activity pattern that was remarkably consistent across stimulation conditions (S2A). This finding is consistent with previously published work examining whole-brain activity changes in animals selectively bred for high versus low habituation rates^37^. Moreover, our data are consistent with a model in which our pharmacogenetic perturbations lead to broad impacts on brain activity. We propose that habituation-influencing manipulations may impact an animal’s internal state, resulting in baseline brain activity changes that are observable via activity mapping, but that manifest at the behavioral level as habituation deficits. Taken together our results support a model in which multiple circuit mechanisms regulated by parallel molecular-genetic pathways cooperate to drive habituation learning *in vivo*.

## Materials and Methods

### Resource Availability

Additional information and requests for resources and reagents reported in this manuscript should be directed to the Lead Contact, Michael Granato.

All data reported in this paper will be shared by the lead contact upon request. All original code is available in this paper’s supplemental information. Any additional information required to reanalyze the data reported in this paper is available from the lead contact upon request.

### Experimental Model and Subject Details

All animal protocols were approved by the University of Pennsylvania Institutional Animal Care and Use Committee (IACUC).

*hip14*^*p*174^, *pappaa*^*p*170^, *cacna2d3*^*sa*16189^, and *ap2s1*^*p*172^ mutants were maintained in the TLF background. *kcna1a*^*p*410^ was maintained in the WIK background. Among these, *cacna2d3*^*sa*16189^ is homozygous viable, and crosses were performed between heterozygous carriers and homozygous mutants to obtain clutches of 50% heterozygous, and 50% homozygous mutant offspring. All other crosses were established between heterozygous carriers.

Pharmacogenetic behavior testing was performed on day 5 as previously described^7^. Stimulation and fixation for MAP-mapping analysis was performed on day 6 to standardize with the reference brain utilized for registration, as previously described^30^.

Clutches were enriched prior to behavior testing or MAP-mapping by selecting for animals based on the exaggerated spontaneous movement phenotype (*kcna1a*^*p*410^) or swim bladder defects (*pappaa*^*p*170^, *hip14*^*p*174^). To enrich for mutant animals in the absence of such phenotypes, *ap2s1* clutches were subjected to live genotyping on Day 3 as previously described^40^. After genotyping, mutant and sibling animals were mixed together in 10cm petri dishes and tested on Day 5 (pharmacogenetic behavior analysis) or stimulated and fixed on Day 6 (MAP-mapping).

For all experiments, behavior was performed and analyzed blind to genotype; genotyping was performed after behavior testing and/or imaging, and mutant animals were compared to siblings from the same clutches.

### Pharmacogenetic Behavior Testing

A 200x (100mM) stock of MK-801 (Sigma M107) was prepared by dissolving a new vial of 25mg of MK-801 powder in 750ul of 100% DMSO. The stock solution was then further dissolved in E3 to a final concentration of 500uM (0.5% DMSO final concentration). MK-801 was applied to a 10cm petri dish containing n=45 5 dpf larvae 30 minutes prior to the first presentation of baseline acoustic stimuli. Control larvae received 0.5% DMSO in E3. The 200x stock of MK-801 was freeze-thawed a maximum of one time and then disposed of.

A 200x (10mM) stock of Strychnine (Sigma S0532) was prepared by dissolving 33.4mg of Strychnine powder in 10mL of 100% DMSO. The stock solution was then further dissolved in E3 to a final concentration of 50uM (0.5% DMSO final concentration). Strychnine was applied to a 10cm petri dish containing n=45 5 dpf larvae 15 minutes prior to the first presentation of baseline acoustic stimuli. Control larvae received 0.5% DMSO in E3. The 200x stock of Strychnine was frozen at −20. We observed no reduction in the effectiveness of our Strychnine stock solution on WT animals over the course of several months of testing.

A 630x (63mM) stock of Butaclamol (Sigma D033) was prepared by dissolving 25mg of Butaclamol in 1mL of DMSO. The stock solution was further dissolved in E3, and DMSO supplemented to a final concentration of 0.5% DMSO, 100uM Butaclamol. Butaclamol was applied to a 10cm petri dish containing n=45 5dpf larvae 30 minutes prior to the presentation of baseline acoustic stimuli. Control larvae received 0.5% DMSO in E3. The 630x stock of Butaclamol was freeze-thawed a maximum of one time and then disposed of.

All larvae were acclimated to testing room conditions (light, temperature, etc.) for 30 minutes prior to the application of pharmacological agents. Assays for habituation of the acoustic startle response (ASR) were performed on 5 dpf larvae arrayed in a 36-well dish, fabricated by laser-cutting a 6×6 grid of holes into an acrylic sheet, and affixing it to an uncut sheet of the same dimensions using acrylic glue. The dish was mounted on a vibrational exciter (4810; Brüel and Kjaer, Norcross, GA) via an aluminum rod. Acoustic stimuli (2ms duration, 1000Hz waveforms) were delivered during the baseline phase of the assay with an interstimulus interval (ISI) of 40 seconds. During the habituation phase, stimuli were presented with a 3-second ISI.

### MAP-mapping

Larvae were acclimated to testing room conditions (light, temperature, etc.) in a 10cm petri dish with n=45 6 dpf larvae for 30 minutes prior to transfer to the testing arena. Following acclimation, 25 larvae were transferred from the petri dish to a cell strainer with 40um pores (Neta Scientific 431750) nested inside a 6cm petri dish, submerged in E3. The entire cell strainer was then removed and immediately submerged in a 4cm petri dish glued to a circular acrylic base, affixed via a titanium arm to the vibrational exciter (4810; Brüel and Kjaer, Norcross, GA). Larvae were acclimated to the testing arena for 30 minutes with no stimuli. “No Stimuli” runs then proceeded with 17 minutes of additional run time. “Non-Habituating Stimuli” runs proceeded with 10 35.1dB acoustic stimuli with a 90-second ISI, followed by 2 minutes of rest. “Habituating Stimuli” runs proceeded with 180 35.1dB acoustic stimuli with 5-second ISI followed by 2 minutes of rest. Immediately following the completion of the behavior testing protocol, the cell strainer was removed from the testing arena and dropped into a 6-well dish (VWR 10861-554) containing 4% PFA in 1x PBT (1x PBS + 0.25% TritonX100). After 2 minutes, cell strainers were transferred to a second 6-well dish containing cold 4% PFA in 1x PBS, and incubated at 4 degrees overnight. Next, the immunostaining procedure described in^30^ was carried out as described with the following modifications: immediately after washing PFA, larvae were bleached for approximately 12 minutes in 1.5% hydrogen peroxide; 0.5% KOH; larvae were then washed twice (quickly) and then once for 5 minutes in PBT; larvae were then incubated in 150mM of Tris-HCl pH 9.0 for 5 minutes at room temperature, followed by 15 minutes at 70 degrees Celcius. Following immunostaining, all larvae for a single experiment were mounted in 1.5% low-melt agarose (Lonza Bioscience 50101) in a 50mm petri dish with a 30mm diameter glass bottom (Mattek P50G-1.5-30-F). Confocal images were acquired using a 20x objective lens on a Zeiss LSM880 confocal microscope using Zen Software. The “tiles” function was used to acquire and stitch together two images of each brain (one centered on the rostral and one on the caudal portion of the head).

### Behavior Analysis

Behavior videos were background subtracted by computing a max projection of the entire image series using FIJI. Max projections were then subtracted from each image within the series using FIJI. Subtracted image series were tracked using FLOTE software as previously described^7,8^. In the case of Strychnine-treated larvae, the previously described “accordion-like” shape^41^ of the SLC response precluded acceptable tracking via FLOTE. Therefore, behavioral responses were manually scored blind to genotype as SLCs or No-Response by isolating the 17ms of video following the delivery of the acoustic pulse, and scoring body bends within this interval as SLCs.

% Habituation was quantified by the following formula: [% Habituation = (1-[response frequency Stimuli 45-54] ÷ [response frequency baseline])*100].

### Quantification and Statistical Analysis

Computation of means, SD, SE, and data set normality were performed using GraphPad Prism. Effects of each drug condition were assessed using Two-Way ANOVA with Sidak’s Multiple Comparisons Test.

For MAP-mapping, image registration and positive and negative significant delta median signals in each brain region across mutant vs. sibling and drug vs. DMSO controls were calculated using the standard MAP-mapping pipeline as described in^30^.

For cluster analysis, positive and negative significant delta median signals were imported into R^42,43^. Signal for each experimental replicate was normalized according to the highest absolute value in that condition, such that the highest magnitude signal for each condition was either −1 or 1. Distances were calculated using the Canberra method, which disregards data when both conditions have a value of 0; this prevented overestimation of similarity between conditions in which many brain regions had zero signal. The factoextra package^44^ was used to visualize distances and clusters.

For pairwise plots, normalized negative signal in a given brain region was subtracted from normalized positive signal to obtain a single signal value for that region. We then averaged signal values for each condition (either drug or mutant allele) over all replicate data sets and behavioral stimulation paradigms. After confirming via cluster analysis that these average values captured the general patterns of similarity observed among individual replicates, we then plotted pairwise comparisons between conditions. Module maps (Figure 7B-D) were created using a modified version of the ZBrainAnalysisOfMAPMaps function^30^. Colors were adjusted in Adobe Illustrator.

## AUTHOR CONTRIBUTIONS

Conceptualization, Writing, Review, and Editing: J.C.N., H.M.S. and M.G. Investigation and Writing-Original Draft: J.C.N. Formal Analysis: J.C.N. and H.M.S. Resources: M.G. Funding Acquisition: J.C.N. and M.G.

## ACKNOWLEDGEMENTS

The authors would like to thank the Granato lab members, as well as Dr. Owen Randlett and Dr. Roshan Jain for technical advice and feedback on our findings. The authors would also like to acknowledge Drs. Roshan Jain, Marc Wolman and Nik Santistevan for sharing *ap2s1*, *pappaa*, and *cacna2d3* mutant zebrafish respectively. The authors would also like to acknowledge the Cell and Developmental Biology Microscopy Core. This work was supported by grants to M.G. (NIH 9R01NS118921) and J.C.N. (1K99NS111736). This article was typeset in Overleaf using the Henriques Lab BioRXiv template with minor modifications.

## Supplementary Figures

**Fig. S1.**
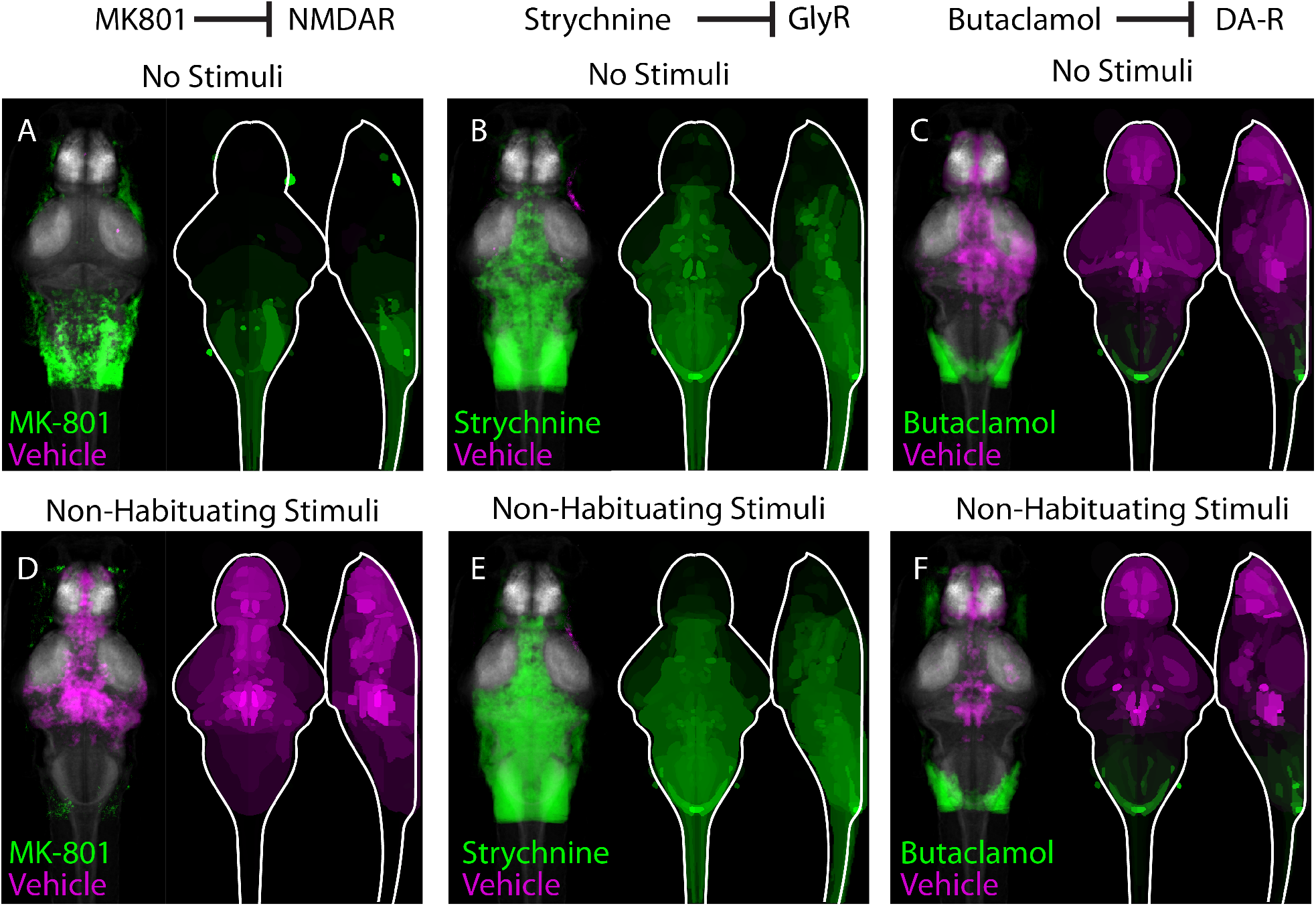
Pharmacological inhibitors of habituation learning induce distinct patterns of brain activity changes. (**A-C**) Regions upregulated by the specified drug treatment under “No Stimulus” conditions are indicated in green; regions downregulated are indicated in magenta. (**D-F**) Regions upregulated by the specified drug treatment under “Non-Habituating Stimuli” conditions are indicated in green; regions downregulated are indicated in magenta. In all images, the left panel is a summed z-projection of the whole-brain activity changes. The right panel is a z-projection and x-projection of the analyzed MAP-map. Molecular targets of pharmacological agents are indicated with diagrams above each column. Note that the patterns of neuronal activity induced a given pharmacological agent are relatively consistent across stimulation condition (i.e. “no stimuli”, vs. “non-habituation stimuli”, vs “habituating stimuli” in Figure 1K-M). Moreover, although all pharmacological agents reduce habituation learning, patterns of neuronal activity are highly dissimilar between individual pharmacological treatments.

**Fig. S2.**
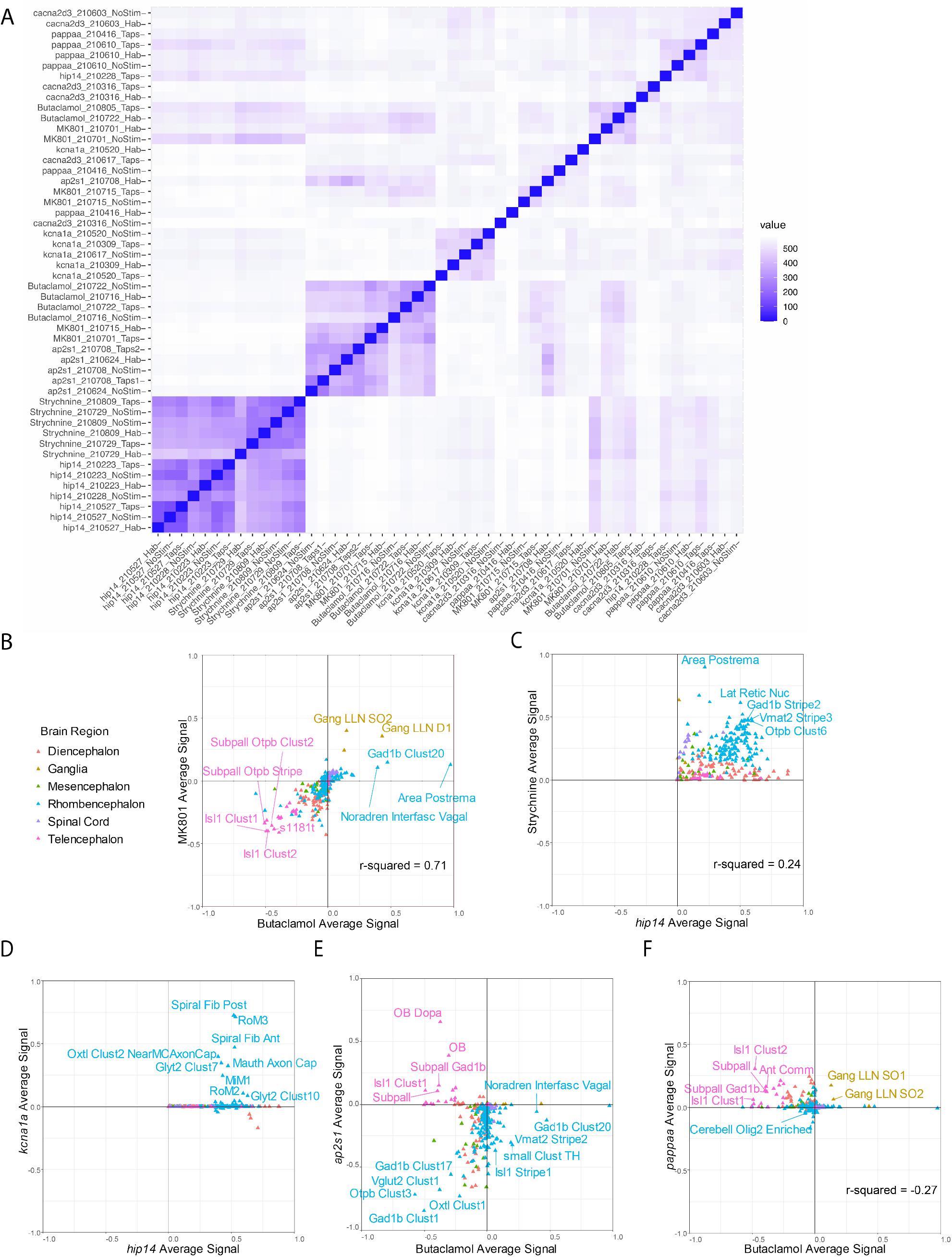
Cluster analysis identifies regions of interest. (**A**) Heat map indicating replicability across stimulus conditions for each mutant and drug condition. Note an intermingled cluster containing Butaclamol and MK-801. (**B-E**) Plots of pairwise comparisons between drugs and or genotypes. Color legend in (**B**) applies to all. R-square values are indicated when p<0.05. (**B**) Plot indicating positive correlation between MK-801 and Butaclamol signal changes. (**C**) Plot indicating a weak positive correlation between *hip14* and Strychnine signal changes. (**D**) Plot showing correlated changes in the rhombencephalon between *hip14* and *kcna1a*. (**E**) Plot showing 3 populations in Butaclamol vs. *ap2s1* changes. Largely telencephalic regions, upregulated by *ap2s1* and downregulated by Butaclamol; largely rhombencephalic regions, upregulated by Butaclamol and downregulated by *ap2s1*; and a large number of regions downregulated by both manipulations. (**F**) Plot showing signal changes in *pappaa* as compared to Butaclamol. Multiple telencephalic as well as diencephalic regions to a lesser degree, are anti-correlated (up-regulated in *pappaa* but downregulated in Butaclamol). Brain region abbreviations in (**B**): s1181t = Telencephalon - S1181t Cluster, Gang LLN SO2 = Ganglia - Lateral Line Neuromast SO2, Gang LLN D1 = Ganglia - Lateral Line Neuromast D1, Noradren Interfasc Vagal = Rhombencephalon - Noradrendergic neurons of the Interfascicular and Vagal areas. Brain region abbreviations in (**C**): Lat Retic Nuc = Rhombencephalon - Lateral Reticular Nucleus. Brain region abbreviations in (**D**): Spiral Fib Post and Ant = Rhombencephalon - Spiral Fiber Neuron Posterior and Anterior clusters, Mauth Axon Cap = Rhombencephalon - Mauthner Cell Axon Cap. Brain region abbreviations in (**E**): DO = Telencephalon - Olfactory bulb dopaminergic neuron areas, OB = Telencephalon - Olfactory Bulb, Subpall Gad1b = Telencephalon - Subpallial Gad1b cluster, Subpall = Telencephalon – Subpallium, Oxtl Clust 1 = Rhombencephalon - Oxtl Cluster 1 Sparse, Noradren Interfasc Vagal = Rhombencephalon - Noradrendergic neurons of the Interfascicular and Vagal areas, TH-C = Rhombencephalon - Small cluster of TH stained neurons. Brain region abbreviations in (**F**): Ant Comm = Telencephalon - Anterior Commisure, Cerebell Olig2 Enriched = Rhombencephalon - Olig2 enriched areas in cerebellum, Gang LLN SO1 and SO2 = Ganglia - Lateral Line Neuromast SO1 and SO2.

## Supplementary Tables

**Supplementary Table S1: Raw values for signal change in each ROI within each mutant and drug condition across 2-3 replicates.**

**Supplementary Table S2: Normalized values for signal change in each ROI within each mutant and drug condition across 2-3 replicates. These values were used for cluster analysis.**

